# Brainstem astrocytes regulate breathing and may affect arousal state in rats

**DOI:** 10.1101/2023.09.28.559604

**Authors:** Mitchell Bishop, Shahriar SheikhBahei

**Author notes:** Corresponding author: Shahriar SheikhBahaei, PhD; Neuron-Glia Signaling and Circuits Unit, National Institute of Neurological Disorders and Stroke (NINDS), National Institutes of Health (NIH), Bethesda, MD 20892, USA.

## Abstract

Variations in arousal levels can impact respiratory patterns. However, whether changes in breathing behaviors can influence arousal state is not fully understood. In this study, we investigated the role of astrocytes in the preBötzinger complex (preBötC) in modulating arousal states via breathing in adult conscious rats. Using viral vector tools, we selectively interfered with astrocytic signaling in the preBötC. Rats with inhibited astrocytic signaling exhibited slower breathing rates and behaviors indicative of a calmer state, whereas enhanced purinergic signaling in preBötC astrocytes led to faster breathing and heightened arousal. Our findings reveal a key role for astrocyte-mediated mechanism in the preBötC that influences both respiratory behaviors and higher-order brain functions like arousal, suggesting a bidirectional link between breathing behaviors and mental states.

**Highlights:**

- In this study, we used molecular approaches to interfere with vesicular signaling in preBötC astrocytes.
- We showed that inhibiting vesicular release mechanisms in preBötC astrocytes are associated with calm behaviors: lower breathing and sigh rates, and longer sniffing time.
- We showed that enhancing vesicular release mechanisms in preBötC astrocytes are associated with more aroused behaviors: higher breathing and sigh rates as well as shorter sniffing time.

## Introduction

Arousal state is defined as the overall alertness at both physical and mental levels [1]. Variations in activities of neurons releasing norepinephrine (NE) in the locus coeruleus (LC) are correlated with the changes in arousal states in mammals [2]–[4]. These LC NE neurons project to the cortex, hippocampus, hypothalamus, amygdala, brainstem, and cerebellum [4], and accordingly affect arousal, memory, and attention [5]. Experimental data show that changes in tonic or phasic firing rate of LC neurons are linked to changes in arousal state [4], [6]–[10].

On the other hand, animals in higher arousal states exhibit increased breathing rate [11],

[12] and decreased regularity of breathing. It is proposed that LC might play a role in this top-down response, as NE has been shown to directly regulate the breathing rate [13]. However, rapid or irregular breathing increases vigilance and arousal [14], suggesting the existence of a bottom-up control of arousal state by breathing [15]. Indeed, recently it was shown that a sub-population of neurons expressing Cdh9/Dbx1 transcription factors in the preBötzinger complex (preBötC, a brainstem center that generate the inspiratory rhythm of breathing [16]) projects to LC and affects arousal state [14]. However, it remains unknown whether breathing behaviors can directly modulate higher order brain function, including arousal.

We have previously shown that preBötC astrocytes can modulate breathing behaviors in conscious behaving rats [17], accordingly, we used viral vector tools to interfere with signaling mechanisms of astrocytes intermingled with the preBötC neurons. We show that rats with slower breathing rates exhibit behaviors associated with calmer state when compared to rats with faster respiratory rates. By modulation of activities of circuits like preBötC, astrocytes can have wide ranging effects [18] and may modulate higher-order brain functions.

## Material and Methods

### Animals

All experiments were performed on adult male Sprague-Dawley rats (250-270g) in accordance with the National Institutes of Health Guide for the Care and Use of Laboratory Animals with project approval from the respective Institutional Animal Care and Use Committees. Animals were housed in a temperature-controlled facility on normal light/dark cycles (12h:12h, lights on at 0700 hours). Tap water and rodent chow were given *ad libitum*.

### Molecular approached to interfere with preBötzinger complex astrocytic signaling mechanisms

To block astrocytic signaling, we used an adenoviral vector (AVV) expressing light chain of Tetanus Toxin (TeLC) under control of an astrocyte specific enhanced GFAP promoter (AVV-sGFAP-GFP-TeLC; 1.3x1012 particles/mL,) [17], [19]. TeLC blocks vesicular exocytosis via degradation of SNARE proteins. Efficiency of TeLC in interfering with vesicular release mechanisms of astrocytes has been described previously [17], [19]–[21]. We also ‘activated’ preBötC astrocytes through stimulation of G_q-_coupled signaling pathway as described before [17]. This was achieved through AVV expressing Gq-coupled Designer Receptor Exclusively Activated by Designer Drug (DREADD_Gq_) fused with green fluorescing protein (GFP) under control of an enhanced GFAP promoter (AVV-sGFAP-DREADD_Gq_-1.3x1012 particles/mL,) [17]. It was previously shown that the expression of DREADD_Gq_ via our AVV-sGFAP-DREADD_Gq_ was constitutively active in the preBötC astrocytes even in absence of ligand, which is typically used in DREADD experiments [17]. We used AVV-sGFAP-CatCh-GFP as a control [17], [20], [21].

### In vivo viral gene transfer

Adult male rats (250–270 g) were anesthetized with a mixture of ketamine/ medetomidine [60 mg kg^−1^/250 μg kg^−1^, intramuscular (i.m.)] and placed in a stereotaxic frame with bregma 5 mm below lambda. PreBötC areas were targeted bilaterally by as described before [17], [21]. Briefly, viral vectors (see above) were delivered via a single microinjection (0.20–0.25 μl) per side at 0.9 mm rostral; 2 mm lateral; and 2.7 mm ventral from the *calamus scriptorium*. After the microinjections, the wound was sutured, and anesthesia was reversed with atipamezole (1 mg kg^−1^, intramuscular). For postoperative analgesia, the animals received buprenorphine (0.05 mg^−1^ kg^−1^ per day, subcutaneous) for 3 days. No complications were observed after the surgery and the animals gained weight normally.

### Measurement of respiratory behaviors in behaving rats

Rodent respiratory activity was measured using whole-body plethysmography. After 5 – 7 days of recovery from viral injections, conscious animals were placed in the Plexiglas chamber (∼2.5L) which was flushed with room air, 22-24 °C, at a rate of ∼2.1 L min^-1^. Levels of O_2_ and CO_2_ in the chamber were monitored by a fast-response O_2_/CO_2_ analyzer (ML206, AD Instruments). We considered the plethysmography chamber as a novel environment and exposure of the animal to this environment then served to induce heightened arousal states [11], [22]. The breathing was recorded during the first 40 minutes after exposure to the plethysmography chamber. All experiments were performed at the same time of day (between 11:00 and 15:00) to account for circadian changes in base level physiology. Data were acquired with Power1401 interface, transferred to *Spike2* software (CED), and analyzed offline.

### Histology and immunohistochemistry

At the end of the experiments, the rats were given an anesthetic overdose (pentobarbitone sodium, 200 mg kg^−1^, intraperitoneal), perfused transcardially with 4% paraformaldehyde in 0.1 M phosphate buffer (pH 7.4), and post-fixed in the same solution for 4–5 days at 4 °C. After cryoprotection in 30% sucrose, serial transverse sections (30–40 μm) of the medulla oblongata were cut using a freezing microtome. One every sixth free-floating tissue sections were incubated with chicken anti-GFP (1:250; Aves Labs, Cat. GFP-1020), rabbit anti-GFAP (1:1000; DAKO, Cat. z-0334), and/or goat anti-ChAT antibody (1:200, EMD Millipore, Cat. AB144P; to confirm location of preBötC) overnight at 4 °C as described before [23]. The sections were subsequently incubated in specific secondary antibodies conjugated to the fluorescent probes (each 1:250; Life Science Technologies) for 1 h at room temperature. Images were taken with a confocal microscope (Zeiss LSM 510).

### Breathing data analysis

Plethysmography data were analyzed as described before [17], [24], [25]. Briefly, the respiratory cycle duration was calculated for each respiratory cycle for the first 40 minutes that the animals were placed in the plethysmography chamber. The respiratory cycles were then averaged in 10-minute intervals and were used to determine the respiratory frequency (number of breaths per minute, *f*_R_). Since it takes about 20-30 minutes for rats to acclimatize to the novel environments, the last 10 minutes were considered as baseline breathing. Tidal volume (V_T_, dept of breathing) and minute ventilation (V_E_ = *f*_R_ x V_T_) were measured as described before [17], [25]. Sighs were defined as peaks with amplitudes two times greater than the average amplitude of the previous four cycles and a respiratory cycle duration 50% greater than the average of the previous four cycles and confirmed manually as described before [17], [24], [25] and reported as the numbers per hour (hr^-1^) The frequency distribution of the instantaneous rate of respiratory-related events (including signing) in the 30-min assay period was analyzed and plotted after extracting high frequency sniffing. The most common breathing rate in each group defined as the dominant breathing frequency (i.e. the mode of the frequency histogram) [11]. High-frequency sniffing was defined as breaths with respiratory rate between 250 and 650 breath min^-1^ [26] and analyzed in the first 10 minutes that an animal was placed in the chamber. Analyzed data were exported to Prism 9.0 software (GraphPad Software Inc., RRID: SCR_002798) where it was reported as averages ± standard error of mean (SEM). For statistical analysis, we used paired or unpaired *t* test, one-way ANOVA, or simple regression analysis as appropriate.

## Results

To affect breathing behaviors in rats, we changed signaling between astrocytes and neurons in the preBötzinger complex (preBötC) by bilateral virally driven expression of either the Gq-coupled designer receptor exclusively activated by designer drugs (DREADD_Gq_) or the light chain of tetanus toxin (TeLC) in astrocytes of the preBötC in adult rats (Figure 1). We used AVV to express CatCh-EGFP in the preBötC astrocytes in the control group [17]. When exposed to a novel environment (i.e., plethysmography chamber), rats expressing the control transgene showed increase in breathing rate by 106 ± 10 min^-1^ (p < 0.001, paired *t* test; *n* = 6) compared to the baseline breathing rate (see method) (Figure 2A&B). When rats that are transduced to express TeLC in preBötC astrocytes exposed to the novel environment, the breathing activity was increased by 77 ± 7 min^-1^ (p < 0.001, paired *t* test; *n* = 6) (Figure 2 A&B). As reported before in the steady-state condition [17], breathing rate in rats expressing TeLC in preBötC astrocytes stayed lower compared to control animals throughout the experiment (Figure 2A). On the other hand, expression of DREADD_Gq_ in preBötC astrocytes increased aroused breathing by 84 ± 3 min^-1^ when compared to that of in the control group (p < 0.001, paired *t* test; *n* = 6; Figure 2B). Compared to the control group, breathing rate started slightly higher and stayed higher in these animals, so the changes in breathing rate were smaller compared to that of control animals (Figure 2A). Since there were differences in baseline and aroused breathing rate between the groups, we then measured the slope of changes in the breathing rate over the time in each group. Simple linear regression analysis suggested that the slope of changes in breathing rate over the first 40 minutes of exposure to the plethysmography chamber were different (F [DFn, DFd] = 2.586 [2, 84]; *p* = 0.081). These data suggest that the rats transduced to express TeLC or DREADD_Gq_ in preBötC astrocytes showed no reduction of the respiratory response when exposed to a novel environment.

**Figure 1.**
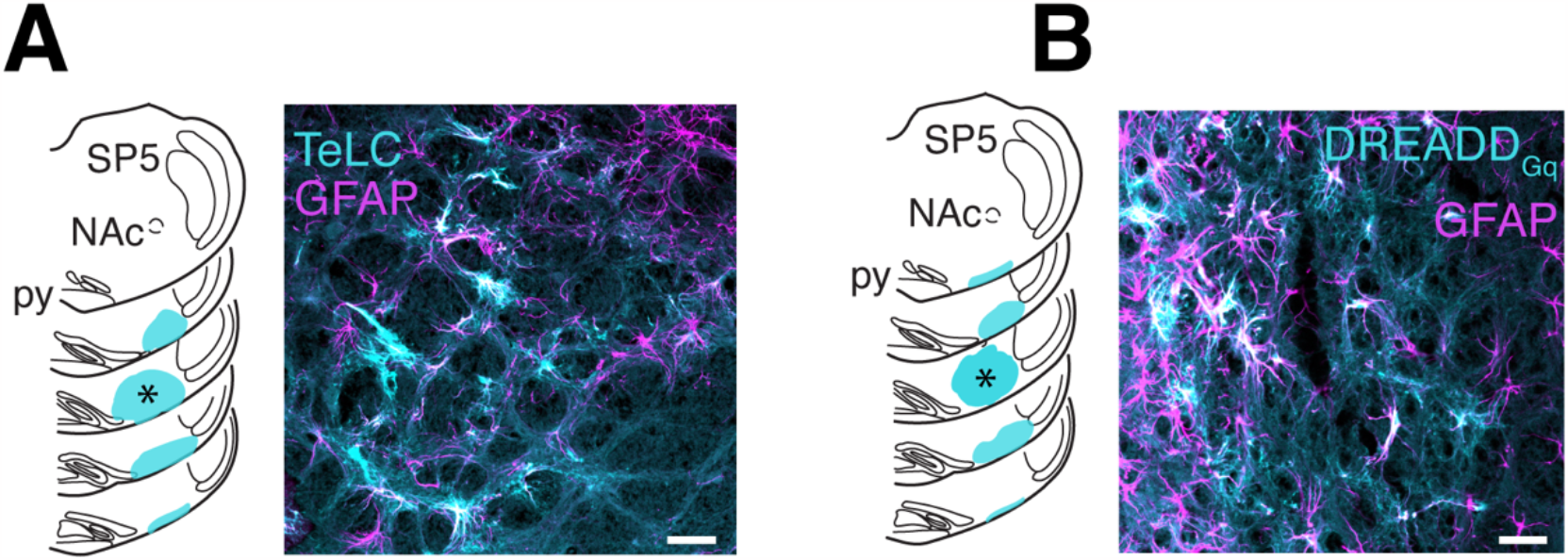
Viral-mediated expression of TeLC and DREADD_Gq_ in preBötC astrocytes. Schematic illustration of the spatial extent of tetanus toxin light chain (TeLC; **A**, *left*) and Gq-coupled designer receptor exclusively activated by designer drugs (DREADD_Gq_; **B**, *left*) expressions following bilateral viral targeting of astrocytes residing within the preBötzinger complex (preBötC) regions of the brainstem ventrolateral medulla. The expression profile (colored regions) for each transgene in the brainstems of all the experimental animals was reconstructed histologically from serial coronal sections. Schematic sections correspond to (from top) -12.1, -12.4, -12.8, -13.1, and -13.4 mm from obex and the star (*) represent location of the preBötC. py, pyramid. NAc, nucleus ambiguus. Sp5, spinal trigeminal nucleus. Confocal images from preBötC region illustrating the expression of TeLC (**A**, *right*) and DREADD_Gq_ (**B**, *right*) in astrocytes residing within the preBötC. Astrocytes expressing the transgenes (cyan) were identified by immunostaining with astrocyte specific marker GFAP (magenta). Scale bars: 50 μm.

**Figure 2.**
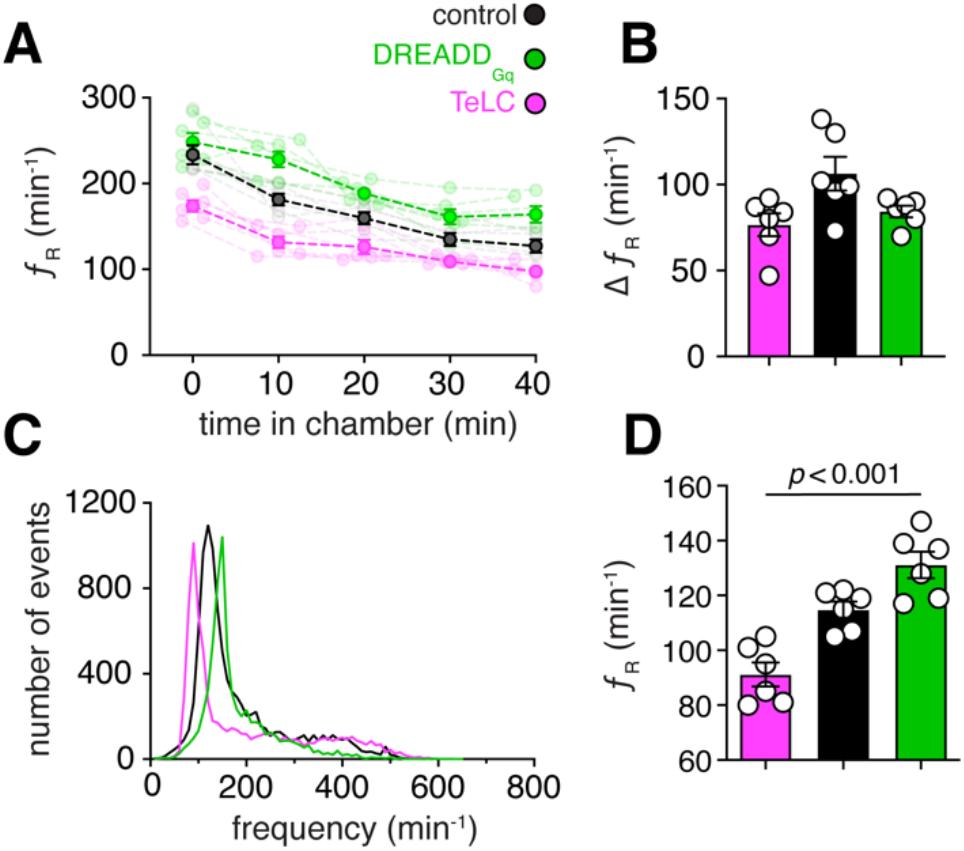
modulation of aroused-breathing frequency by preBötC astrocytes. **(A)** Changes in breathing frequency (*f*_R_) during the first 40 min in the plethysmograph in rats expressing DREADD_Gq_, TeLC, and control transgene in preBötC astrocytes (n = 6 animals per group). Data are shown as individual (lighter colored lines) and mean values ± SEM (darker colored lines). **(B) S**ummary data illustrating the difference between breathing rate (Δ*f*_R_) at 0 and 40 minutes after the animal is exposed to the plethysmography chamber. **(C)** Group data showing a representative diagram of effects of TeLC or DREADD_Gq_ expression in preBötC astrocytes on frequency distribution detected in first 40-min of a conscious adult rat is in the plethysmography chamber. Expression of DREADD_Gq_ shifted the dominant *f*_R_ (peak) to a higher frequency (right shift) while expression of TeLC shifted the dominant frequency to the left. **(D)** Group data showing the effect of TeLC or DREADD_Gq_ expression in preBötC region on dominant *f*_R_ (peaks found in **C**) in conscious rats. Data in **B** and **D** are presented as mean ± SEM. Each data point represent measurement obtained in one animal. *p* values – one way ANOVA test.

The dominant breathing frequency was found to be higher in rats transduce to express DREADD_Gq_ (131 ± 5 min^-1^) and lower in rats expressing TeLC (91 ± 4 min^-1^) in preBötC astrocytes when compared to that of in control animals (131 ± 5 min^-1^; one-way ANOVA, *p* < 0.001, F (2, 15) = 23.66; *n* = 6 per group) (Figure 2C&D). We also measured tidal volume and minute ventilation and consistent with previous data [17], [27], expression of TeLC or DREADD_Gq_ in the preBötC astrocytes did not affect tidal volume or minute ventilation in rats during the experiment (data not shown).

Since sigh (augmented breath) frequency (*f*_S_) has been proposed to be related to the arousal level, we next analyzed *f*_S_. Rats expressing TeLC in preBötC astrocytes had lower *f*_S_ (24 hr^-1^) and rats transduced to express DREADD_Gq_ had higher *f*_S_ (39 hr^-1^) after exposure to the novel environment compared to the control rats (29 hr^-1^; one-way ANOVA, *p* < 0.001, F (2, 15) = 19; *n* = 6 per group) (Figure 3A).

**Figure 3.**
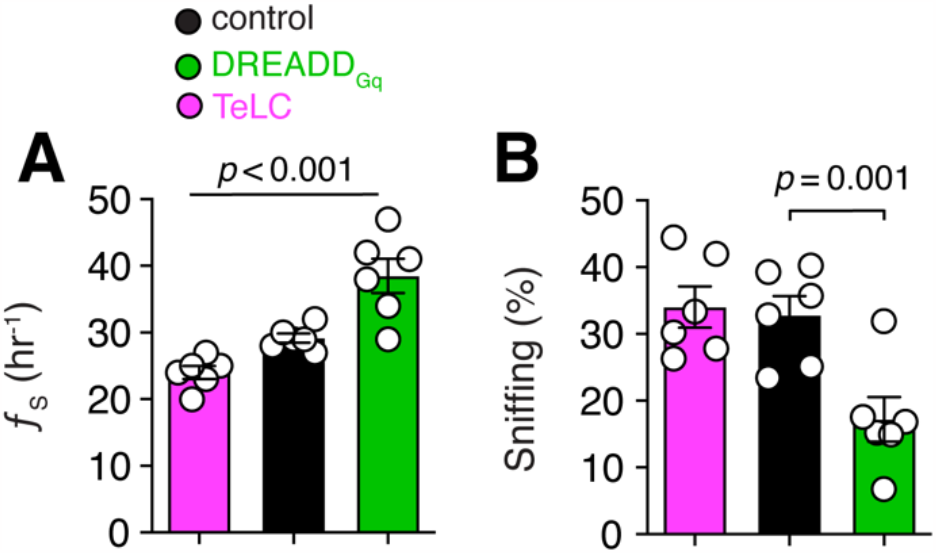
**(A)** Summary data showing frequency of sighs (*f*_S_) in the first 40-min that a conscious rat is in the plethysmography chamber. Rats transduced to express TeLC in the preBötC displayed lower *f*_S_, while rats expressing DREADD_Gq_ in the preBötC astrocytes showed higher *f*_S_. **(B)** Group data depict the time spent by the animals at high-frequency sniffing, expressed as % of total time (5-min epochs). Rats expressing DREADDGq showed lower sniffing time, while rats expressing TeLC, or control (CatCh-EGFP), engaged in sniffing behavior about the same amount of time. Data are presented as mean ± SEM. Each data point represent measurement obtained in one animal. *p* values – one way ANOVA (**A**) and unpaired *t* test (**B**).

*f*_R_ and *f*_S_ data suggested that rats transduced to express TeLC might be calmer, and rats transduced to express DREADD_Gq_ might have higher arousal state compared to the control group. We then measured high frequency sniffing times as it is linked to aroused or calm behaviors in rodents [11], [28]. Animals transduced to express TeLC in preBötC astrocytes had similar sniffing time to the control group (34 ± 3 % vs. 33 ± 3 % in control; p = 0.7, unpaired *t* test, *n* = 6 per group) (Figure 3B). Interestingly, rats expressing DREADD_Gq_ in the preBötC astrocytes showed shorter sniffing time (17 ± 3 % vs. 33 ± 3 % in control; p = 0.001, unpaired *t* test, *n* = 6 per group) (Figure 3B) compared to the control group. These data suggest that expression of DREADD_Gq_ in preBötC astrocytes is linked with less exploratory high frequency sniffing behaviors.

## Discussion

This study focused on the preBötC, the fundamental brainstem respiratory rhythm-generating circuit, and investigated whether changes in breathing rate can affect arousal state in conscious rats. It has been shown that preBötC astrocytes can regulate activities of preBötC neurons [17], [21]; therefore, we used molecular approaches to block (via bilateral viral expression of TeLC) or stimulate preBötC astroglial vesicular release of gliotransmitters (via bilateral viral expression of DREADD_Gq_) to decrease or increase breathing rate, respectively. TeLC efficiently blocks Ca^2+^-dependent vesicular exocytosis in preBötC astrocytes [17], [19], and expression of TeLC in preBötC astrocytes is shown to reduce the *resting* breathing rate (*f*_R_), frequency of periodic sighs (*f*_S_), and impair respiratory responses to physiological challenges [17].

In our study, the expression of TeLC or DREADD_Gq_ in astrocytes was restricted to the preBötC (Figure 1), therefore, in these rats, we have affected breathing behaviors that are known to be generated by the cells in the preBötC region, such as rhythm and pattern of breathing [16], [29] and sighs [30]. Similarly, our data suggest that rats transduced to express TeLC in preBötC have lower *f*_R_ and *f*_S_ during exposure to a novel environment (Figures 2D & 3A). On the other hand, previous data suggest that expression of DREADD_Gq_ in preBötC astrocytes increases *f*_R_ and *f*_S_ at rest [17]. Consistent with these data, our result also showed that rats transduced to express DREADD_Gq_ in preBötC astrocytes have higher *f*_R_ and *f*_S_ during exposure to novel environment as well (Figures 2D & 3A). We have shown before that interfering with normal signaling mechanisms of preBötC astrocytes in rats only affects *f*_R_, but not V_T_ and V_E_, at room air with intact peripheral chemosensors [17], [25], [27], [31]. Consistent with these data, we also observed changes *f*_R_ in rats transduced to express TeLC or DREADD_Gq_ in preBötC astrocytes when exposed to novel environment. Since some neurons of the preBötC project to other regions in the brain [32], [33] [14], it is plausible that we have also affected neural activities that are either originating at the level of the preBötC or affected by the preBötC, including changes in arousal state [14].

It was shown previously that our AVV that expresses DREADD_Gq_ in preBötC astrocytes is constitutively active as expression of this transgene led to a higher level of phospholipase C (PLC) activity, higher rate of spontaneous vesicular fusion, and increased tonic release of ATP [17]. Therefore, no ligand was necessary to activate DREADD_Gq_ and consequently, there is no added stress because of intraperitoneal injection of ligand to the animal. We exploit this property of our DREADD_Gq_ construct to evaluate if sustained activation of preBötC astrocytes has an impact on the arousal state. In addition to higher *f*_R_ and *f*_S_, rats transduced to express DREADD_Gq_ in preBötC astrocytes displayed lower sniffing and showed no reduction of the respiratory response to the chamber. Lower sniffing time has been shown to be associated with higher arousal state as the animal is less likely to explore the environment [11], [34]. On the other hand, rats transduced to express TeLC in preBötC astrocytes showed respiratory behaviors that are representative of calmer behaviors [11].

In this study, we focused on the respiratory behaviors associated with arousal states. Other rodents’ behaviors, such as grooming, are also linked to calmer state [14], and future experiments are needed to characterize other behaviors associated with calmer state in rats transduced to express TeLC or DREADD_Gq_ in preBötC astrocytes. Together with our data, these results will enhance our comprehension of the relationship between respiratory patterns and cognitive performance. A further constraint of the present research is the absence of data on the potential modulatory effects of anxiolytic agents on the respiratory alterations observed in rats transduced to express DREADD_Gq_ in preBötC astrocytes. Nevertheless, the data obtained in the present study indicate that normal function of the astrocytes may be important to mount the appropriate physiological response during mild aroused states. These data further support the hypothesis that the role of astrocytes may become particularly important in conditions where homeostatic regulation of brain circuits is critical to support behavioral and physiological demands.

**Table 1.** linear regression analysis.

|  | control | TeLC | DREADD<sub>Gq</sub> |
| --- | --- | --- | --- |
| ***Best-fit values*** |  |  |  |
| Slope | -2.59 | -1.758 | -2.357 |
| R squared | 0.7483 | 0.7085 | 0.6941 |
| *Y-intercept* | 219.1 | 162.8 | 245.2 |
| *X-intercept* | 84.61 | 92.61 | 104 |
| *1/slope* | -0.3861 | -0.5687 | -0.4243 |
| **Std. Error** |  |  |  |
| Slope | 0.2839 | 0.2131 | 0.2957 |
| *Y-intercept* | 6.954 | 5.221 | 7.243 |

## Acknowledgements

We thank NINDS Light Microscopy Core for technical assistances. We are grateful to Drs. Jeffrey Smith and Alexander Gourine for support and mentorship and thank Dr. Sergey Kasparov for providing the viral vectors. This work was supported by the Intramural Research Program of the NIH, NINDS and NIMH (ZIA NS009420 to SSB).

## Notes

Conflict of interest: The authors declare no competing financial interests.

### Competing Interest Statement

The authors have declared no competing interest.

